# Self-beneficial belief updating as a coping mechanism for stress-induced negative affect

**DOI:** 10.1101/2020.12.02.408096

**Authors:** Nora Czekalla, Janine Baumann, David S. Stolz, Annalina V. Mayer, Johanna F. Voges, Lena Rademacher, Frieder M. Paulus, Sören Krach, Laura Müller-Pinzler

## Abstract

Being confronted with social-evaluative stress elicits a physiological and a psychological stress response. This calls for regulatory processes to manage negative affect and maintain self-related optimistic beliefs. The aim of the current study was to investigate the affect-regulating potential of self-related belief updating after exposure to social-evaluative stress, in comparison to non-social physical stress or no stress. We assessed self-related belief updating using trial-by-trial performance feedback and described the updating behavior in a mechanistic way using computational modeling. We found that social-evaluative stress was accompanied by an increase in cortisol and negative affect which was related to a shift in self-related belief updating towards the positive direction. This self-beneficial belief updating, which was absent after physical stress or control, was associated with a better recovery from stress-induced negative affect. This indicates that enhanced integration of positive self-related feedback can act as a coping strategy to deal with social-evaluative stress.

## Introduction

Human beings strive to be accepted by others and to maintain a positive social image^1^. Thus, social evaluation from others can pose a threat to our social image, eliciting a stress response in our body^2–4^. This initiates various physiological processes^5^ and is associated with negative affective consequences, like anxiety or embarrassment^6–8^. Social evaluation, however, is fundamental to self-related learning processes, as it gives one the opportunity to integrate the feedback we receive from others and update the beliefs about ourselves accordingly^9,10^. Biases in how we process self-related feedback, i.e. whether we focus more on negative or positive feedback, impact our affective reactions^11,12^ and, in the case of self-serving processing, may function as a coping strategy^11^. While (social) stress is a risk factor for many psychiatric conditions^13^, successful coping is an important factor in maintaining mental health^14^. In the current study we implemented a computational modeling approach to investigate the coping mechanism of self-beneficial belief updating after social-evaluative stress and tested whether shifted information processing after stress predicts recovery from stress-induced negative affect.

When we receive feedback regarding ourselves, information processing and belief updating is shaped by self-relevant motivations^15^, especially the motivation to maintain optimistic beliefs about the self^16^. Many studies have demonstrated that the process of self-related belief updating is biased in favor of positive information, i.e. self-related beliefs are updated more strongly when feedback is better than expected^17–20^. However, updating biases towards negative feedback have been reported in performance contexts^21,22^, which indicates that the context of learning, type of feedback and prior assumptions are important factors when explaining self-related belief updating biases.

While there are only relatively few studies on the effects of stress on self-related belief updating, various studies on reward processing and non-self-related feedback processing have shown that stress is an influencing factor in this regard. One key mechanism for feedback-based learning is the prediction error signal, indicating the difference between a predicted and an actual outcome^23,24^, which is being minimized by updating beliefs during learning. This signal is generated by dopaminergic neurons of the ventral striatum^25^, which might be particularly important for the stress-induced modulation of prediction error signals as the dopamine system is sensitive to stress^26,27^. However, these effects depend on the type, intensity and schedule of the stress exposure^28^, which might also explain heterogeneous effects of stress on reward processing and feedback-based learning. Research on declarative memory has shown that timing of stress matters^29^, which seems to be important for feedback-based learning as well^30^. Initially, acute stress (e.g. a threat of a shock during learning), mainly characterized by a rapid sympathetic response, impairs feedback-based learning of reward^31^. Neurally, acute stress attenuates the response to reward in the striatum and orbitofrontal cortex^32,33^ and enhances the striatal response to aversive feedback^34^. Accordingly, under acute stress self-related belief updating is more strongly driven by unfavorable feedback, i.e. the learning bias in favor of positive information (optimism bias) usually found in self-related belief updating is absent^35^. The opposite effects are reported when learning takes place with a delay to stress (e.g. after a public speech), a phase mainly characterized by an increase of cortisol^29^. Here, non-self-related feedback processing is more strongly driven by stimuli signaling reward and possibly associated with stress-induced cortisol change^36^ while learning from negative feedback is decreased, potentially linked to cortisol levels before learning^37^. On the neural systems level, stress recovery is associated with increased striatal responses to rewarding feedback at 50 min after stress^30,38^. Moreover, specifically individuals with low striatal reward reactivity showed an association of recent life stress with lower positive affect, which makes striatal reactivity a potential factor of successful stress coping^39^.

According to classic appraisal theories of stress^40^, different strategies such as seeking social support, positive revaluation or acceptance are helpful in coping with stress-induced negative affect^40–42^. In the context of social-evaluative stress a self-protection strategy is to view oneself in a positive light, i.e. emphasizing the own desirability, focusing on own successes and attributing failure externally^43^. This strategy has also been successful in alleviating stress-induced negative affect following a performance situation^11,44^. Generally, an optimistic way of processing self-related feedback has been associated with better mental health^45,46^. On the contrary, processing self-related feedback in a more negative way may result in negative beliefs about the self^47^ and ultimately lead to lower self-esteem or depressive symptomatology. Studies on self-related belief updating in individuals with depression suggest that information processing is distorted in a negative direction^48^ and that coping strategies for situations of social-evaluative stress are less readily available in these patients^49^.

In the present study, we aim to investigate the specific effects of social-evaluative stress on self-related belief updating and the propensity to engage into self-beneficial belief updating after social-evaluative stress as compared to non-social physical stress. By means of two well validated and highly reliable paradigms, the Trier Social Stress Test^3^ (public speech) and the Cold Pressor Test^50^, as well as a no stress control condition, we directly manipulated levels of social-evaluative stress in a between-groups design. After stress manipulation we assessed participants’ self-related belief updating behavior with the learning of own performance (LOOP) task^21^, in which participants form beliefs about their abilities in novel behavioral domains. We then used participants’ learning bias from positive and negative feedback to predict their recovery from stress-induced negative affect. We found that social but not physical stress shifted subsequent self-related belief updating in a more self-beneficial direction which predicted better recovery from negative affect. We elaborate on the relationship between stress, self-related belief updating and affect regulation in healthy participants and discuss the potential of our findings for a better understanding of maladaptive self-related belief systems in psychiatric conditions such as depression.

## Results

After exposure to social-evaluative stress (SOC, Trier Social Stress Test), non-social, physical stress (PHY, Cold Pressor Test) or a no stress control condition (CON, reading) participants performed the LOOP task^21^, which was covered as a measure of cognitive estimation skills (see Fig. 1). The central idea of the LOOP task is to create a performance context and provide manipulated positive or negative feedback in comparatively neutral domains in which people have only vague prior assumptions. By this means, individuals form a concept about their own abilities over the course of the experiment. In a previous study, we showed that this process of self-related belief updating can be described best by a computational prediction error learning model (adapted from Rescorla and Wagner^51^) with two separate learning parameters for positive and negative prediction errors^21^. During the LOOP task, participants were asked to answer estimation questions in two different estimation domains (e.g. estimating the weight of animals and the height of buildings) and received manipulated performance feedback implying a rather good performance in one category and a rather bad performance in the other one (high vs. low ability condition). In the beginning of each trial participants saw a cue indicating the estimation category and had to rate their expected performance for the upcoming estimation question in this category. A manipulated feedback on their estimation performance in relation to an alleged reference group was presented afterwards. Saliva cortisol as well as negative affect, including perceived stress, embarrassment, anger, and frustration, were assessed several times during the experiment. Pre-stress baseline measures (T1_AFF/CORT_) were taken after a ten-minute-period of rest in the beginning of the session. Post-stress negative affect was rated immediately after the stress exposure or control task (T2_AFF_) to calculate the mean change of negative affect (ΔAFF). Post-stress cortisol samples were taken after another 10-minute period of rest (T2_CORT_) to calculate the mean cortisol change (ΔCORT). After performing the LOOP task, saliva samples and negative affect were again obtained (T3_AFF/CORT_) (for a detailed description see methods).

**Fig. 1.**
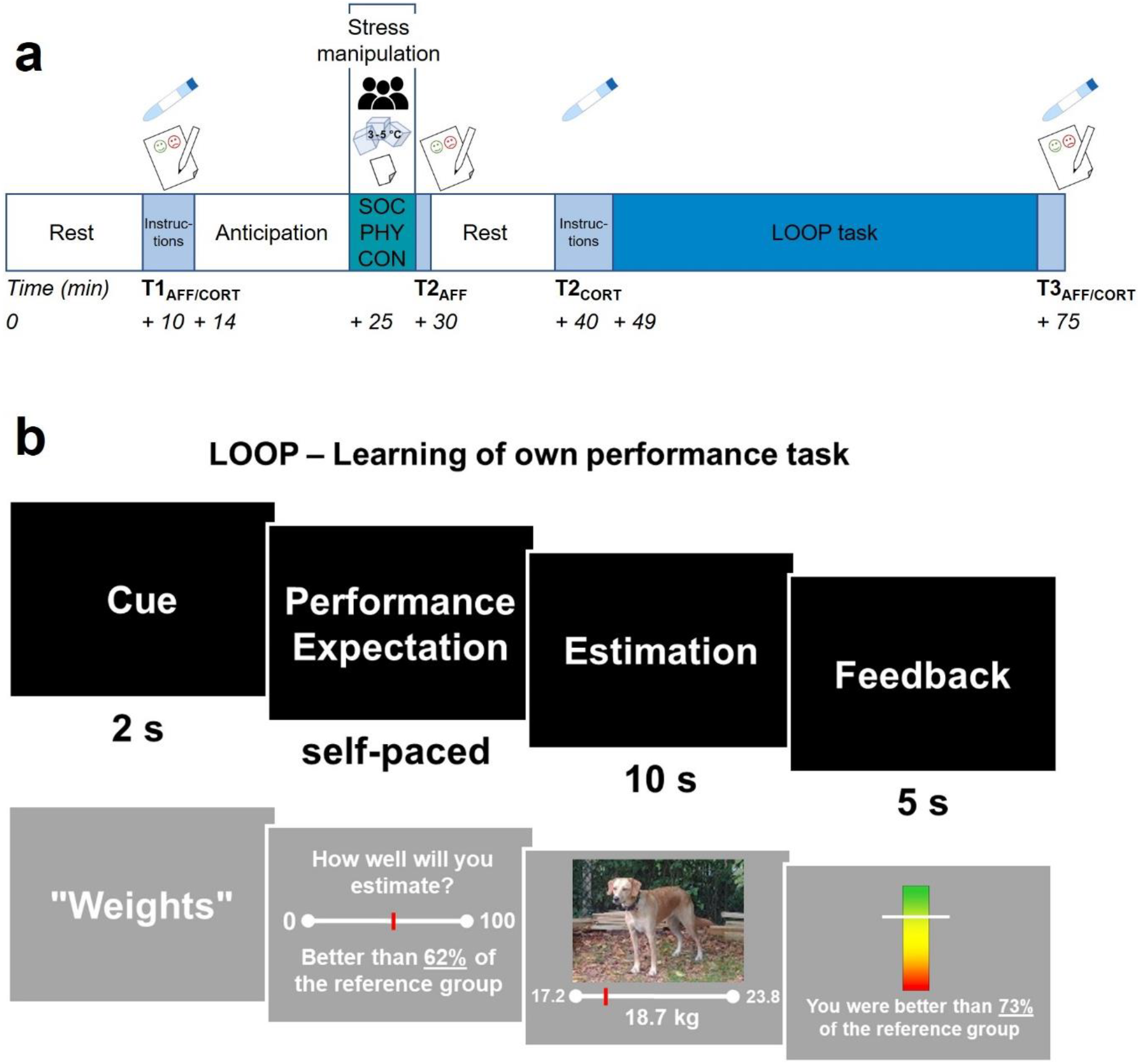
**(a)** Experimental timeline and procedure. SOC: social-evaluative stress group (public speech [audience icon], n = 29), PHY: physical stress group (Cold Pressor Test [ice cubes icon], n = 30), CON: no stress control group (reading task [paper icon], n = 30), salivette icon: saliva collection for cortisol determination; paper pencil icon: rating of negative affect including perceived stress, embarrassment, anger, and frustration. **(b)** Sequence of one trial. 1. Cue: display of the upcoming estimation category associated with a high or low ability condition, 2. Performance expectation rating, 3. Estimation question, 4. Performance feedback. Figure adapted from Müller-Pinzler et al.^21^.

### Cortisol response and negative affect

#### Cortisol

The stress manipulation was effective and social-evaluative stress, as well as physical stress, led to a stronger increases of cortisol levels from baseline T1_CORT_ to post-stress T2_CORT_ than in the no stress control group (Scheirer-Ray-Hare test controlled for time of the day [TIME]: main effect factor Stress group H_2_ = 18.9, *p* < .001, post-hoc Dunn-Bonferroni-Tests for factor Stress group: SOC vs. CON: z = −4.29, *p* < .001; PHY vs. CON: z = −2.76, *p* = .018). There was no statistically significant difference between the two stress groups (SOC vs. PHY: z = 1.56, *p* = .355; baseline cortisol levels did not significantly differ between groups *H*_2_ = 1.74, *p* = .419 controlled for TIME, see Fig. 2a and Supplementary Fig. S1 and Table S1).

**Fig. 2.**
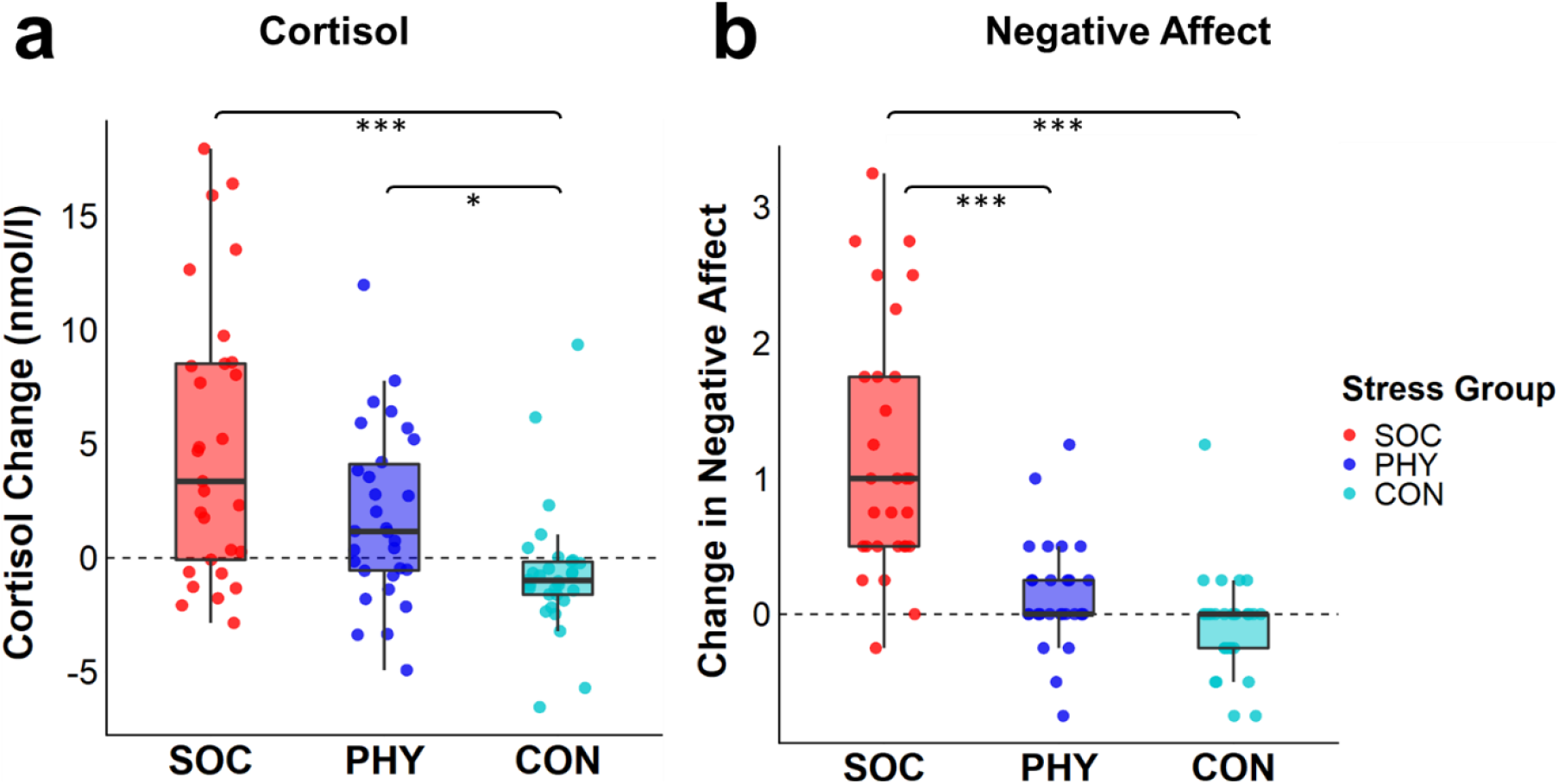
**(a)** Change in saliva cortisol levels after stress induction (post-stress T2_CORT_ - baseline T1_CORT_), **(b)** Change in negative affect (post-stress T2_AFF_ - baseline T1_AFF_), SOC = Social-evaluative stress group, PHY = Physical stress group, CON = no stress control group. Line inside box: median, lower/upper box hinges: 25^th^ and 75^th^ percentile, lower/upper box whiskers: smallest/largest value within 1.5 x inter-quartile range from hinges, \**p* < .05, \*\*\**p* < 001.

#### Negative affect

Mean negative affect increased significantly after social-evaluative stress but not after physical stress compared to the control group (Kruskal Wallis test: H_2_ = 43.9, *p* < .001, post-hoc Dunn-Bonferroni-tests: SOC vs. CON (*z* = −6.45) *p* < .001, PHY vs. CON (*z* = −1.88) *p* = .182, SOC vs. PHY (*z* = − 4.59) *p* < .001; baseline negative affect did not significantly differ between groups (H_2_ = 3.2, *p* = .201; see Fig. 2b and Supplementary Fig. S2).

### Forming self-related beliefs over time

In a model free behavior analysis we replicated previous findings regarding the LOOP task which indicates self-related belief updating in response to the feedback^21^. Over the time of 30 trials, participants adapted their performance expectation ratings (EXP) towards the positive and negative feedback of the two ability conditions, i.e. they updated their self-related beliefs (Fig. 3a, significant factor Ability condition high vs. low *t*_86_ = 8.52, *p* < .001, significant Trial x Ability condition interaction *t*_5156_ = 32.72, *p* < .001). Social-evaluative stress modulated self-related belief updating over time, i.e. performance expectation ratings became increasingly higher compared to physical stress or no stress (Trial x Ability condition x Stress group split into the contrasts social [SOC] vs. non-social [PHY, CON] and the orthogonal contrast PHY vs. CON: interaction for contrast SOC vs. [PHY, CON]; *t*_5156_ = 4.01, *p* < .001). In the physical stress group performance expectation ratings were even more negative over time than in the no stress control condition (Trial x Ability condition x Contrast PHY vs. CON *t*_5156_ = −2.15, *p* = .031; see Supplementary Table S2).

**Fig. 3.**
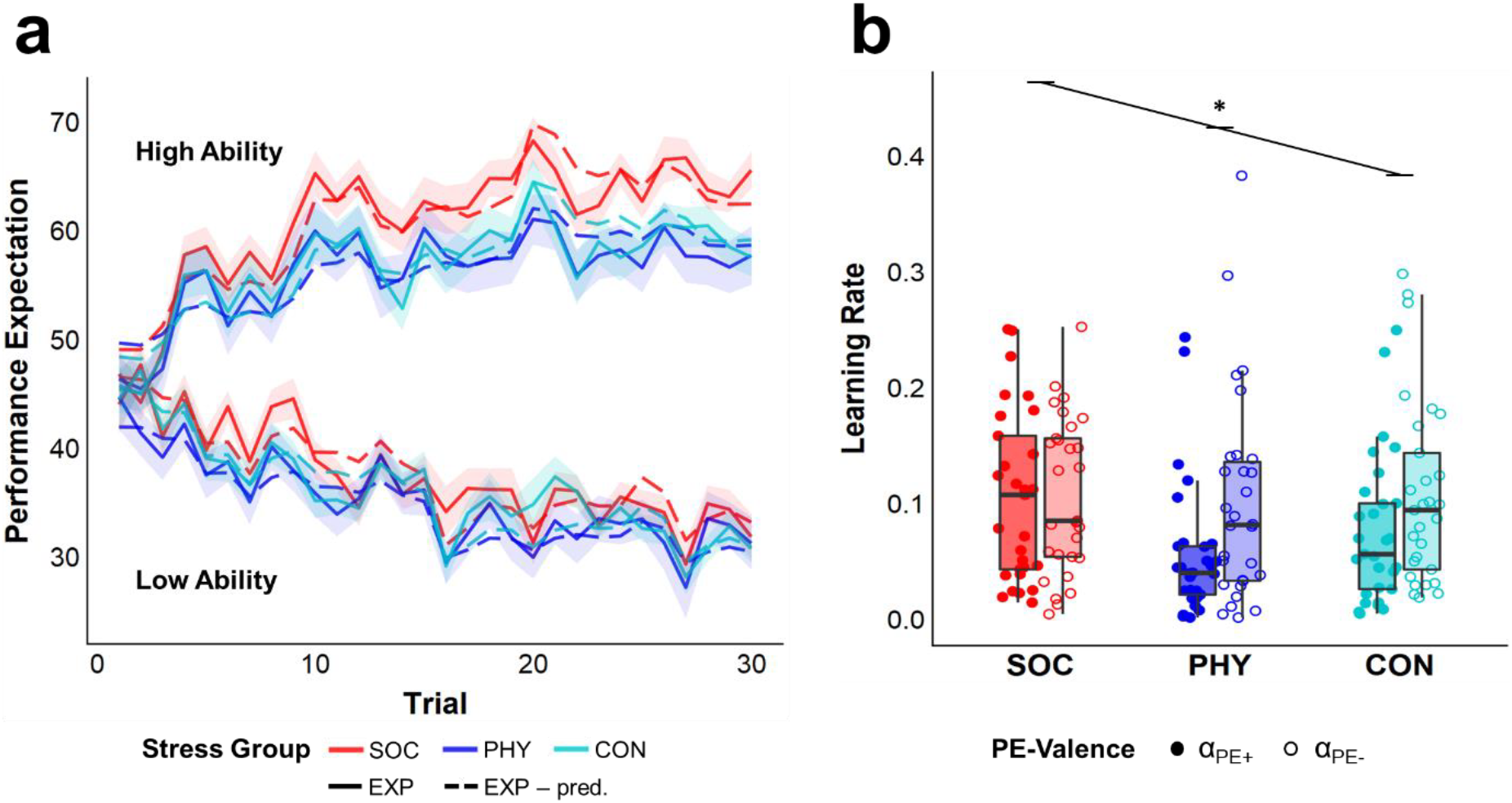
**(a)** Performance expectation ratings (EXP, solid line) and performance expectations predicted by the winning model (EXP – pred., dashed line) over the time course of 30 trials. Ratings and predicted values were averaged across participants separately for the two ability conditions and the three experimental groups. Shaded areas represent the standard errors of the expectation ratings for each trial. **(b)** Learning rates derived from the Valence Model. A significant interaction effect (*) of PE-Valence x Stress group (SOC = social-evaluative stress, PHY = physical stress, CON = no stress control) indicates that a bias towards increased updating in response to negative prediction errors (α_PE−_) in contrast to positive prediction errors (α_PE+_) is absent in the social-evaluative stress group.

### Model selection for computational models of learning behavior

To capture the updating of the performance expectation ratings over time in a learning model, a similar model comparison to that of Müller-Pinzler et al.^21^ was performed. All three main models of the model space followed the idea of a Rescorla-Wagner model with one or two learning rates for each participant reflecting the degree to which people weighted prediction errors (PE = Feedback_t_ - EXP_t_) to update their expectation rating (see Fig. 4 and for model descriptions see method section).

**Fig. 4.**
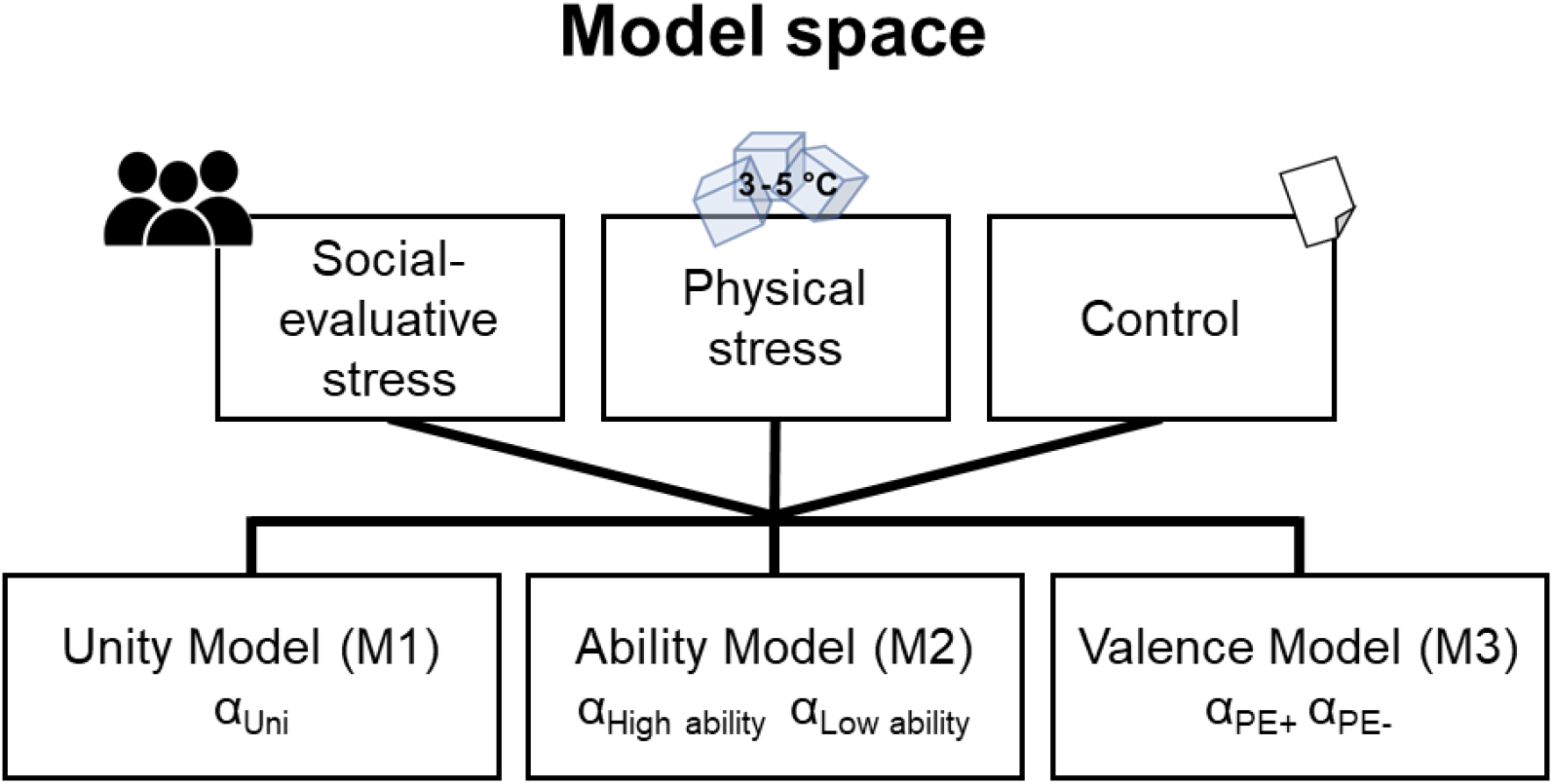
Structure of the model space. α_Uni_ = one learning rate for the whole time course; α_High ability_/ α_Low ability_ = two separate learning rates for the two ability conditions; α_PE+_/ α_PE−_ = two separate learning rates for positive and negative prediction errors; Figure adapted from Müller-Pinzler et al.^21^..

In line with Müller-Pinzler et al.^21^, the Valence Model outperformed all other models in all three groups according to Bayesian Model Selection^52^ (protected exceedance probability for the whole sample *pxp*_total_ > .999, Bayesian omnibus risk *BOR*_total_ < .001 as well as separately for the three groups *pxp*_SOC_ = .985, *BOR*_SOC_ = .019, *pxp*_PHY_ > .999, *BOR*_PHY_ < .001, *pxp*_Control_ > .999, *BOR*_Controll_ < .001; see Table 1 and Supplementary Table S3 for more details on model comparisons). This model, with two separate learning rates for positive PEs (α_PE+_) and negative PEs (α_PE−_) across ability conditions, assumes that learning differs depending on the valence of prediction errors. Learning parameters from the Valence Model were used for further analysis.

**Table 1.**
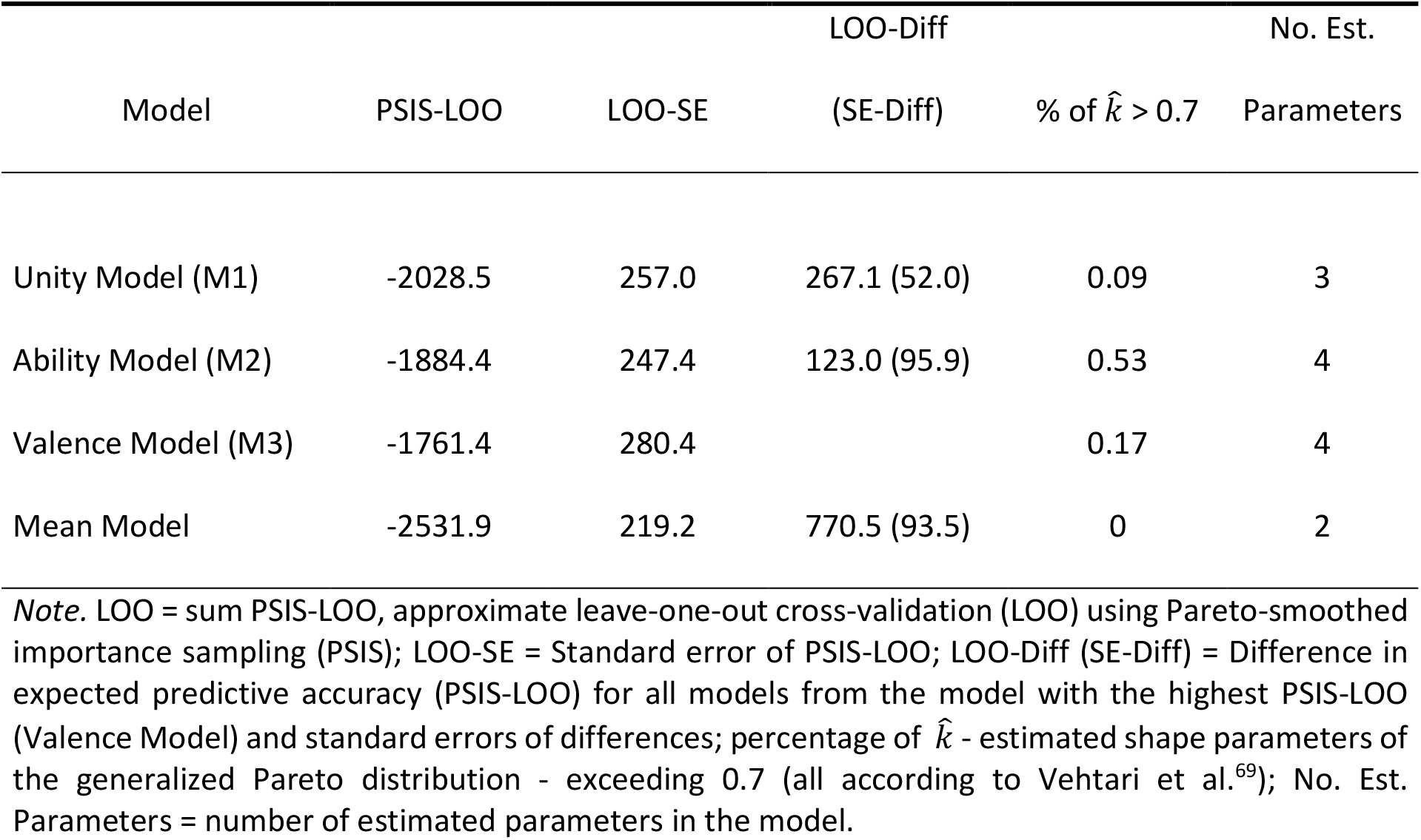
PSIS-LOO Scores for the whole sample

The modeled performance expectations of our winning model predicted the performance expectation ratings on the individual subject level within each ability condition with *R^2^* = 0.33 ± 0.24 (*M* ± *SD*). Repeating the model free analysis with the modeled performance expectations confirmed the results from the original analysis (see Supplementary Table S4).

### Stress and learning parameters

In line with Müller-Pinzler et al.^21^, the physical stress and no stress control group showed a negativity bias in their learning behavior, i.e. a stronger self-related belief updating after negative than positive prediction errors (α_PE+_ vs. α_PE−_ within group comparison for PHY: *W* = 100, *Z* = −2.73, *p* = .005 and CON: *W* = 84, *Z* = −2.89, *p* = .003, Wilcoxon test). This negativity bias was absent after social-evaluative stress (α_PE+_ vs. α_PE−_ within group comparison for SOC: *W* = 193, *Z* = − 0.53, *p* = .609; significant PE-Valence x Contrast SOC vs. [PHY, CON] interaction *b*_VALxSOC_ = 0.114, *t*_85_ = 2.30, *p* = .024, PE-Valence x Contrast PHY vs. CON: *b*_VALxPHY_ = −0.036, *t*_85_ = −0.72, *p* = .471; betas standardized, see Fig. 3b and Supplementary Table S5).

To better capture biased learning behavior, a valence bias score was computed (valence bias score = (α_PE+_ - α_PE−_)/(α_PE+_ + α_PE−_))^21,53,54^, which represents updating after positive compared to negative prediction errors. More positive valence bias scores indicate more self-beneficial belief updating, while negative valence bias scores speak for stronger self-related belief updating after negative feedback.

#### Negative affect predicts subsequent self-beneficial belief updating

We found that a stronger increase in negative affect (T2_AFF_ - T1_AFF_) predicted more self-beneficial belief updating (*ρ*_ΔAFF,BIAS_ = .25, *p* = .019, Spearman correlation for the whole sample, see Fig. 5b). Also, a higher increase in in cortisol levels (saliva samples T2_CORT_ - T1_CORT_) predicted more self-beneficial belief updating (*ρ*_ΔCORT,BIAS|TIME_ = .29, *p* = .006, partial Spearman correlation controlled for TIME for the whole sample, for further information see Supplementary Results and Supplementary Table S6).

**Fig. 5.**
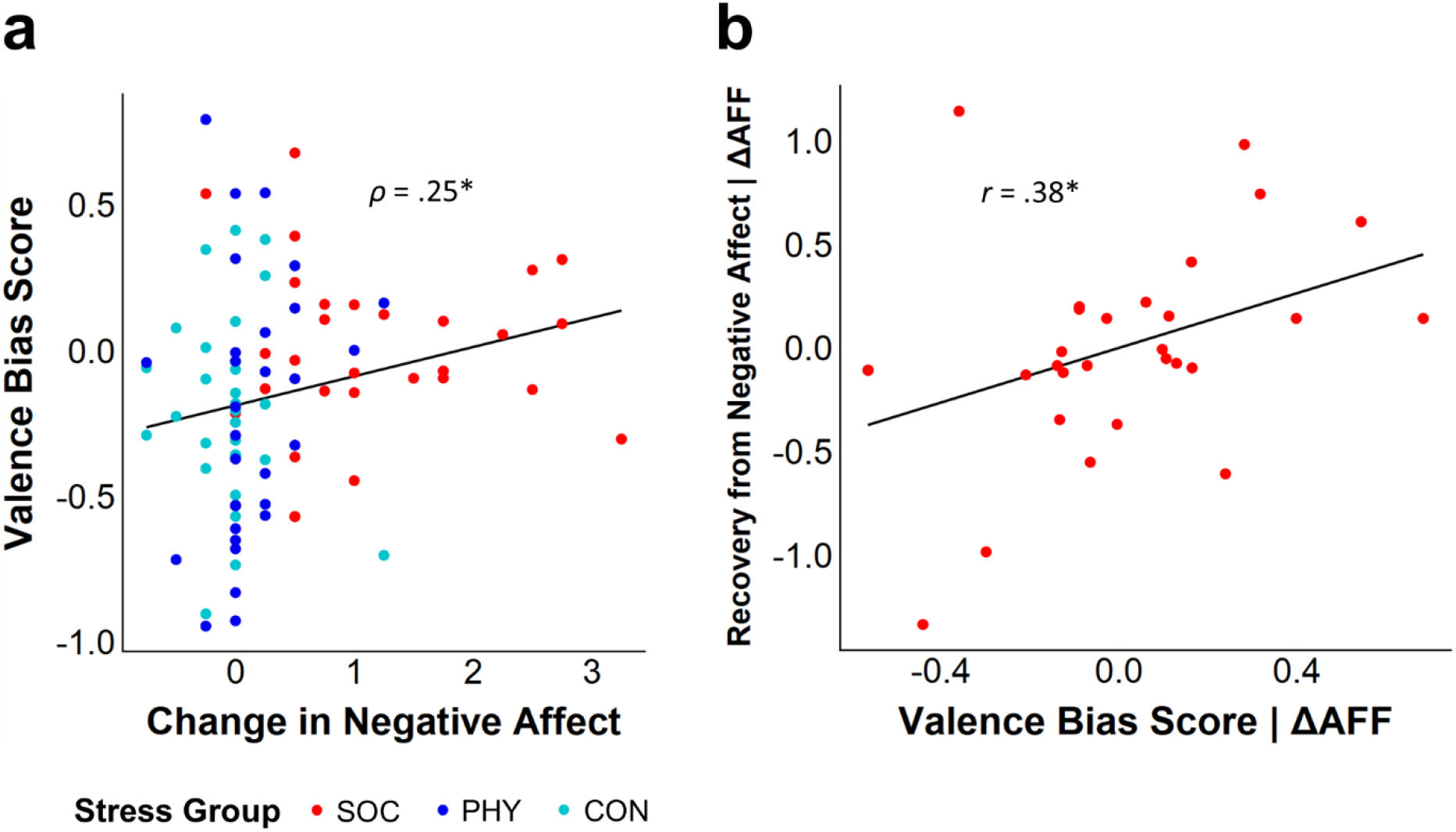
**(a)** Correlation plot of valence bias score ((α_PE+_ - α_PE−_)/(α_PE+_ + α_PE−_)) and stress-induced change in negative affect (ratings T2_AFF_ - T1_AFF_). A stronger affective stress response was associated with more self-beneficial belief updating (higher valence bias score) in the subsequent learning paradigm. Slope of a linear regression model added for better visualization; *ρ* = Spearman’s Rho. **(b)** Partial correlation plot of valence bias score and recovery from negative affect (REC, ratings T2_AFF_ - T3_AFF_) in the subsample of the social-evaluative stress group (n = 29) controlled for the stress-induced change in negative affect (ΔAFF, ratings T2_AFF_ - T1_AFF_). More self-beneficial belief updating (higher valence bias score) is associated with a better recovery from stress-induced negative affect. Slope fit with linear regression model; *r* = Pearson’s *r*; * = *p* < .05.

#### Learning rates and affective recovery

A more positive valence bias score predicted better recovery from stress-induced negative affect during learning (REC, change in negative affect post-stress T2_AFF_ - post-learning T3_AFF_, *ρ*_BIAS,REC|ΔAFF_ = .23, *p* = .043 [partial Spearman correlation for the whole sample controlled for the increase in negative affect (T2_AFF_ - T1_AFF_)]). This supports the idea of self-beneficial belief updating as a coping strategy. Analysis of the social-evaluative stress group only confirmed this effect (Fig. 5b, *r*_BIAS,REC|ΔAFF_ = .38, *p* = .045 [partial Pearson correlation controlled for the increase in negative affect]), for correlations within the three experimental groups see Supplementary Table S7). Further exploratory analysis on the relationship between the valence bias score and the recovery from cortisol (T2_CORT_ −T3_CORT_) did not reveal any significant effect (see Supplementary Table S8).

## Discussion

After being devalued for example at work or school we need to empower ourselves in order to uphold or boost our self-image. Research has shown that the ability to adopt a positive attitude towards oneself after receiving criticism is central to positive affect and good mental health outcomes in the long run^11,46,55^. In the current study we investigated how people apply self-beneficial belief updating during a performance feedback situation as a means to counter their negative affect. Using computational modelling, we provide a mechanistic explanation on how individuals engage in more self-beneficial belief updating after experiencing a threat to their social image and how this shift in social learning of self-related information predicts recovery from stress-induced negative affect.

The positive shift of self-related belief updating after social-evaluative stress, going along with a better recovery from negative affect, fits nicely to the notion of a belief’s own value as recently posited by Bromberg-Martin and Sharot^15^. In their revised framework, general belief updating is not solely driven by external outcomes like rewards or punishments but also by the agent’s motivation to optimize internal states like positive affect^15^. In the present study we show this direct link between self-related belief updating and a change in the affective state indicating that self-related belief updating might be motivated by the wish to uphold or even recover a positive affective state. This is in line with the idea of motivated cognition, i.e. the assumption that cognitive processes like attention, information processing and decision making are not neutral on their own, but are always shaped by needs, feelings and desires of the individual^56^. Especially when processing information that challenges one’s self-image, self-related belief updating is not only informed by the history of previous feedback, as it has often been assumed in classical reinforcement learning tasks, but also by various self-relevant needs and goals^57^. Transferred to the present study, this implies that the motivation to restore an endangered self-image and to regulate one’s affect back to a set point directly impacts self-related information processing. The pattern of an active counter-regulation of negative affect by self-beneficial belief updating can be described as a striving for homeostasis^11^. To better capture the fluctuation of the affective state and its involvement in the trial-by-trial self-related belief updating loop, following the framework by Bromberg-Martin and Sharot^15^, future studies should consider repeated assessments of affective states during the task to predict the empowering potential of shifts in learning on the single trial level.

Since negative self-related beliefs are at the core of psychiatric conditions like depression^47^, this study targets clinically highly relevant processes. Depression is associated with seeking negative feedback which confirms negative self-related beliefs^58^ and seeking negative feedback in combination with a stressful life event can even increase depressive symptoms^59^. Furthermore, depression is associated with a weaker stress recovery mediated by an attentional bias towards negative feedback. Understanding the mechanisms of how people form self-related beliefs in a context mimicking everyday performance settings and linking these to the regulation of negative affect after stress has important implications for understanding the etiology of depressive symptoms. The present study set-up, including a social-evaluative stress induction followed by a social-evaluative performance situation also addresses one of the fundamental fears of individuals with social anxiety: being devalued by others. Since both depression and social anxiety are associated with negatively biased updating behavior in response to self-related feedback^12,21,48,60^, we assume that the affect-regulating and empowering potential of self-beneficial belief updating after social-evaluative stress would be less pronounced in depression or social anxiety and would thereby possibly exacerbate the symptomatology in a self-fulfilling way. Future studies with similar experimental set-ups and clinical samples could examine the relationship between self-beneficial belief updating and affect regulation in more detail and develop potential intervention strategies based on empowering individuals on their way to processing newly incoming information.

Replicating a previous study of ours^21^, self-related belief updating was negatively biased in the control condition in which participants were not exposed to any stress. In the prior study, this negativity bias has been shown to be specific for self-related belief updating in comparison to belief updating about another person^21^. In the present study, we found that after physical stress participants also exhibited a negativity bias in forming self-related beliefs, i.e. participants tended to make greater updates in response to negative prediction errors in contrast to positive prediction errors. The negativity bias stands in contrast to other studies reporting a positivity or optimism bias in feedback-based learning e.g. when receiving feedback about the chance to encounter negative life events^20,61^, about one’s intelligence^17^ or about one’s personality^18,19^ (for a review see^16^). There are several possible explanations for the motivation behind the negativity bias in context of the LOOP task in contrast to the reported positivity biases of other studies which was, however, not the focus of the present study (for a discussion on the negativity bias see^21^). In order to test for the specificity of self-beneficial belief updating after social-evaluative stress, it would be interesting to test if this effect also accounts for experiments that typically yield a positivity bias (e.g. for life events, IQ or personality) in feedback-based learning tasks.

Here, we demonstrated that both, negative affect and cortisol stress responses, go along with a shift in self-related belief updating. It has been shown before that experiencing social emotions (e.g. embarrassment or shame) is related to increased cortisol levels in situations which threaten one’s social image, like the social-evaluative stress induction^6^. Cortisol has been linked to reward processing and feedback-based learning in the *stress triggers additional reward salience - STARS* - model which proposes that stress and the associated release of cortisol modulates the dopamine system, resulting in an increased salience of rewards, thus biasing learning towards rewarding feedback^36,62^. The current results, however, suggest that the quality of stress (here, social vs. physical) might make a difference, and the *STARS* model, based on a rather unspecifically triggered cortisol response, cannot fully explain the present stress effect on self-related belief updating after social but not physical stress. While in our study both measured components of social-evaluative stress, negative affect and the cortisol stress response, were associated with shifts in learning behavior, we cannot rule out that our alteration of the Cold Pressor Test, to remove the social element of the conductor, potentially resulted in some participants terminating the test prematurely. This mode of testing might have been less intense and a more intense physical stress protocol might lead to similar effects on learning behavior. A more detailed recording of negative affect as well as the physiological stress response might help in future studies to better differentiate between different stress qualities and understand specific effects of social-evaluative stress.

To summarize, our results indicate a shift towards more self-beneficial belief updating after social-evaluative but not physical stress. This shift goes along with a better recovery from stress-induced negative affect. Linking self-related belief updating to affect is an important step in understanding biases in self-related learning and its relation to affect regulation. The special feature of the present study was the study-set that allowed to examine a link between negative affect and self-related belief updating. By introducing a performance context with consecutive self-related feedback, corresponding to real-life school or work related performance situations, individuals can form beliefs about their own abilities over time and potentially use this formation process as a means to regulate their affect. With this approach we aimed to increase the ecological validity of the study in order to trigger and investigate motivational processes that might be less relevant in more abstract study settings. Since social evaluation represents a constant stressor in every-day life, the question of an appropriate coping strategy to regulate negative affect is of great importance when handling everyday social situations.

## Materials and Methods

### Participants

Eighty-nine participants recruited at the University of Lübeck Campus were included in the study. Upon appearance, participants were assigned to either a social-evaluative stress group (SOC; *n* = 29, 21 female, aged 18–28 years; *M* = 22.9; *SD* = 2.76), a physical stress group (PHY; *n* = 30, 20 female, aged 19–27 years; *M* = 22.5; *SD* = 1.94) or the control group (CON; *n* = 30, 20 female, aged 18– 32 years; *M* = 22.3; *SD* = 3.00, data of the control group were published before^21^). From the initially recruited *N*=96 subjects, seven had to be excluded – five because they did not believe the cover story and two due to technical problems. All included participants were fluent in German, non-smokers with a body-mass index between 18.5 and 30. They were not diagnosed with acute or chronic psychiatric conditions or diseases affecting the hormone system and did not take psychiatric drugs or medication affecting the hormone system (except hormonal contraceptives). Participants had normal or corrected-to-normal vision and did not study psychology to avoid previous experience with experiments using cover stories. Additional exclusion criteria for participants who underwent the physical stress protocol were cardiovascular diseases, frequent fainting or seizures and current hand injuries. For more details on the sample characteristics see Supplementary Table S9a. All participants gave written informed consent prior to the participation and received monetary compensation for their participation. They were naive to the background of the study during the session and debriefed about the cover story afterwards. The study was conducted in compliance with the ethical guidelines of the American Psychological Association (APA) and was approved by the ethics committee of the University of Lübeck.

### Manipulation procedure

#### Social-evaluative stress

Social-evaluative stress was induced by a public speech similarly to the Trier Social Stress Test^3^. Participants were instructed to prepare a short self-presentation for an application for a scholarship, which had to be presented in front of a selection committee who would allegedly assess the participant’s verbal skills and body language. The selection committee consisted of the experimenter, who was passive during the speech, a second experimenter, who was allegedly responsible for measuring verbal skills, and a passive camera assistant, who pretended to videotape the speech. Before starting the ten-minute preparation period, participants briefly visited the room with the selection committee. After the preparation time was over, participants were asked to come back to this room and present their speech. Talking time was five minutes (*M* = 4.9 min, *SD* = 0.16) with a minimum of three minutes of uninterrupted speech. If the participant finished the speech before the time was over, the second experimenter waited for at least 15 seconds with a motionless face and then asked the participant to continue. If the participant stopped speaking again and the three minutes of free speech had passed, the second experimenter asked standardized questions until the five minutes of talking time were over (“Explain why it is important for you to achieve a good performance.”, “Do you think it is important to improve yourself throughout your life?”, “Do you consider yourself a person who values his/her independence?”). Average social-evaluative stress duration (start subsequent rest period – start speech preparation) was *M* = 16.4 min, *SD* = 1.2.

#### Physical stress

Physical stress was induced by an exposure to ice water according to the Cold Pressor Test protocol^50,63^. Participants were asked to dip their non-dominant hand in cold water (water temperature 3 – 5.5°C = 37.4 – 41.9°F, *M* = 4.26°C, *SD* = 0.50) for as long as possible up to three minutes (duration 48 sec – 3 min, *M* = 2.7 min, *SD* = 0.7). The water was kept in motion with a small electrical pump to prevent the water temperature from rising around the participant’s hand. To control for the procedure of the social-evaluative stress condition, participants visited the room with the cold pressor apparatus first, had a ten-minute preparation period and came back into the room for the stress exposure. During the preparation time, participants were asked to imagine dipping their hands in a freezing cold environment and write down their associations. To make the stress exposure less social, the experimenter was not present in the room but waited in an adjacent room. If the participant took out their hand before the three minutes were over, they had to signal this immediately by ringing a bell. The experimenter could roughly observe the participant in the reflection of the glass door, thus ensuring that she/he dipped the hand into the water. Average physical stress duration (including preparation period) was *M* = 16.2 min, *SD* = 1.5.

#### No stress control condition

In the control condition, participants performed a reading task that was described to them as measuring reading speed. They had ten minutes to rehearse two different texts about applying for a scholarship. Afterwards, they were guided to the other room with nobody present and were asked to measure their reading time, while reading the two texts aloud at a natural speed. Average control duration was *M* = 15.3 min, *SD* = 1.3.

### Manipulation checks

#### Cortisol

Three saliva samples were collected during the experiment for cortisol analysis (see Fig. 1a). The first sample (baseline T1_CORT_) was taken after a 10 min period of rest immediately before starting the instruction for the stress manipulation (mean time between T1_CORT_ and start of the SOC, PHY or CON preparation phase: *M* = 3.7 min, *SD* = 1.4). The post-stress cortisol sample T2_CORT_ was collected after another 10 minutes resting period following the stress manipulation and the last sample (T3_CORT_) was collected after the learning task (*M* = 45.6 min (*SD* = 3.3) post stress). The stress-induced cortisol change (ΔCORT) was determined by subtracting the cortisol levels of T2_CORT_ - T1_CORT_. Saliva was collected with Salivettes (Sarstedt, Nümbrecht, Germany), stored at −30 °C and sent to the bio-psychological lab at TU Dresden, Dresden, Germany for analysis (here stored at −20 °C until analysis). Salivary free cortisol levels were determined using a chemoluminescence immunoassay (IBL International, Hamburg, Germany).

#### Negative affect

We assessed negative affect by means of a short pen and paper questionnaire, covering the emotions embarrassment, anger, frustration, as well as the perceived stress with one rating each. The questionnaires were handed out at baseline (T1_AFF_) as well as at the very end of the experiment (T3_AFF_). The post-stress negative affect was measured immediately after the stress manipulation (T2_AFF_; see Fig. 1). Ratings were averaged for each measurement point to get a composite measure of negative affect (see Supplementary Fig. S2 for separate scores). The change in negative affect after stress (ΔAFF) was determined by subtracting T1 negative affect from T2 (T2_AFF_ - T1_AFF_). The recovery from negative affect (REC) was determined by subtracting T3 negative affect from T2 (T2_AFF_ - T3_AFF_).

### Behavioral task

#### Learning of own performance task

The Learning of own performance (LOOP) task^21^ (Fig. 1b) allows to measure self-related belief updating through trial-by-trial performance expectation ratings and subsequent performance feedback. The task included estimation questions in two different estimation categories (heights of houses and weights of animals) and was presented to the participants as a measure of estimation abilities. To make participants learn about their estimation ability the two estimation categories were paired with manipulated performance feedback implying high ability for one category and low ability for the other (e.g. heights of houses = high ability and weights of animals = low ability, estimation categories were counterbalanced between ability conditions). The assignment of the categories to the ability conditions was independent of the participants’ actual performance and their performance expectation ratings. Thus, participants could learn over the course of the experiment that they were good in one estimation category and rather bad in the other one. Each trial began with a cue displaying the category of the next estimation question followed by a performance expectation rating for this question. Afterwards, the estimation question was presented together with a picture for ten seconds. Continuous response scales below the pictures determined a range of plausible answers for each question, and participants indicated their responses by navigating a pointer on the response scale with a computer mouse. Subsequently, feedback indicating the estimation accuracy as percentiles compared to an alleged reference group of 350 university students was presented for five seconds (e.g. “You are better than 72 % of the reference participants.”). The order of the two estimation categories/ability conditions was intermixed with a maximum of two consecutive trials of the same condition and 30 trials per condition in total. The estimation questions were randomized within the estimation category/ability conditions. A fixed sequence of ability conditions and feedback was presented for all participants. In the low ability condition, feedback was approximately normally distributed around the 35th percentile (SD ≈ 16; range 1–60%) and in the high ability condition around the 65th percentile (SD ≈ 16; range 40–99%). The task started with detailed instructions and three test trials. All stimuli were presented using MATLAB Release 2015b (The MathWorks, Inc.) and the Psychophysics Toolbox^64^.

### Procedure

To minimize noise in the cortisol saliva samples, participants were asked to follow behavioral rules prior to the experimental session. These were in detail: no alcohol on the evening before the experiment and bed rest at about 10 p.m. (ideal case eight hours of sleep); one hour before the session: no sport, no smoking, no drinks containing caffeine or theine, no food (including bonbons and chewing gums) and no juices. Upon arrival at the laboratory, participants read the participant information including the cover story regarding the stress manipulation and the LOOP task. After signing the consent form, they were asked to fill out a questionnaire checking the adherence to the behavioral rules. Participants rested for ten minutes before the baseline measurement, including saliva cortisol and negative affect, was obtained (T1_AFF/CORT_). During the resting period, they filled out a short personality questionnaire (not included in this study). Subsequently, participants of the social and physical stress groups were challenged with a stress protocol while participants of the control group did the control reading task. Directly afterwards, participants rated their affective state (T2_AFF_) followed by another ten minutes resting period, which was terminated with a saliva sampling (T2_CORT_). In the second part of the experiment participants performed the LOOP task. Finally, another cortisol sample and affective ratings were collected (T3_AFF/CORT_). After completing a post-experimental interview, including additional questionnaires, participants were debriefed about the cover story. The experimental sessions were run between 10.00 a.m. - 12.00 p.m., 1.00 - 3.00 p.m. or 3.45 - 5.45 p.m. The allocation to the time slots did not differ between the experimental groups (Pearson’s Chi-squared test *p* = .867, see Supplementary Table S9b). See Fig. 1a for a graphical illustration of the procedure.

### Statistical analysis

#### Stress manipulation

To test whether the stress manipulation was effective, the stress-induced changes in cortisol as well as affect were compared between the three experimental groups. Due to the stress manipulation, the variance of the cortisol and negative affect responses were unequal between the three experimental groups (Levene test *p*s < .05). Since the distributions of the cortisol and affective stress response were skewed in some groups (Lilliefors-corrected Kolmogorov-Smirnov normality test *p*s < .05) non-parametric tests were used. Negative affect responses were compared with the Kruskal-Wallis test, the cortisol response was compared with the Scheirer-Ray-Hare test, an extension of the Kruskal–Wallis test, to control for time of the day (morning vs. noon vs. afternoon, see Procedure). Post-hoc comparisons between the groups were performed with Dunn’s test.

#### Model free analysis of performance expectation ratings

The analysis of the expectation ratings including computational modeling was adapted from Müller-Pinzler et al.^21^. To illustrate basic effects of the expectation ratings, a linear mixed model with the factors Ability condition (high ability vs. low ability), the continuous variable Trial (30 Trials), and Stress group (with the two contrasts SOC vs. [PHY, CON] and PHY vs. CON) as a between subject factor was performed.

#### Computational modeling of learning behavior

The dynamic changes in self-related beliefs, which were measured by the performance expectation ratings in response to the provided performance feedback, were modeled using prediction error delta-rule update equations (adapted from Rescorla-Wagner model^51^). There were three main models of the model space with one or two learning rates modeled separately for each participant (see Fig. 4). The first model (Unity Model) included a single learning rate for the whole time course (EXP_t+1_ = EXP_t_ + α_Uni_ PE_t_). The second model (Ability Model) contained two separate learning rates for the two ability conditions allowing to capture a difference in expectation updating when receiving feedback in a high ability context (α_High ability_) or low ability context (α_Low ability_). The third model (Valence Model) with two separate learning rates for positive PEs (α_PE+_) and negative PEs (α_PE−_) across ability conditions allows to model learning that differs depending on the valence of prediction errors rather than different ability conditions. The three models were compared to a Mean Model with two performance expectations means reflecting the assumption of stable expectations for each ability condition without learning over time. In addition to the learning rates, we fitted two parameters for the initial belief about participant’s performance, separately for both ability conditions (see Table 1).

#### Model fitting

For model fitting we used the RStan package^65^, which uses Markov chain Monte Carlo (MCMC) sampling algorithms. All learning models of the model space were fitted separately for each subject. To sample posterior parameter distributions, a total of 2400 samples were drawn after 1000 burn-in samples (overall 3400 samples; thinned with a factor of 3) in three MCMC chains. Convergence of the MCMC chains to the target distributions was assessed by 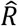 values^66^ for all model parameters. One subject was excluded due to implausible model parameters, i.e. mean learning rate of almost 1, as well as 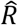 values of 1.1 and low effective sample sizes (*n*_*eff*_, estimates of the effective number of independent draws from the posterior distribution) for some model parameters of the valence model. Otherwise the effective sample sizes were greater than 1000 (>1400 for most parameters). Posterior distributions for all parameters for each of the participants were summarized by their mean resulting in a single parameter value per subject that we used to calculate group statistics.

#### Bayesian model selection and family inference

To select the model that describes the participants’ updating behavior best, we estimated pointwise out-of-sample prediction accuracy for all fitted models separately for each participant by approximating leave-one-out cross-validation (LOO)^67^. To this end, we applied Pareto-smoothed importance sampling (PSIS) using the log-likelihood calculated from the posterior simulations of the parameter values as implemented by Vehtari et al.^67^ (loo R package^68^). Sum PSIS-LOO scores for each model as well as information about 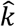 values, the estimated shape parameters of the generalized Pareto distribution, indicating the reliability of the PSIS-LOO estimate, are depicted in Table 1. As summarized in Table 1 very few trials resulted in insufficient parameter values for 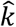 and thus potentially unreliable PSIS-LOO scores (on average 0.20 % of trials per subject with 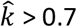). Bayesian model selection on PSIS-LOO scores was performed on the group level accounting for group heterogeneity as described by Stephan et al.^52,69^. This procedure provides the protected exceedance probability for each model (*pxp*), indicating how likely a given model has a higher probability explaining the data than all other models, as well as the Bayesian omnibus risk (*BOR*), the posterior probability that model frequencies for all models are all equal to each other^69^. Additionally, difference scores of PSIS-LOO for all models in contrast to the winning model were computed, which can be interpreted as a simple ‘fixed-effect’ model comparison^67^ (see Table 1).

#### Posterior predictive checks

To test whether the predicted values of the winning model could capture the variance in the performance expectation ratings a regression analysis (EXP ~ pred. values) was performed for each subject separately for the two ability conditions. R-squared statistic was determined and averaged. In addition, the model free analysis of the expectation ratings was repeated with the predicted values of the winning model to assess if the predicted data captured the effects that were present in the data of the expectation ratings.

#### Analysis of learning parameters

Learning rates for positive (α_PE+_) and negative prediction errors (α_PE−_, factor PE-Valence) were compared between the three groups in a linear mixed model with the factors PE-Valence and group (split into the contrasts SOC vs. [PHY, CON] and PHY vs. CON). Additional post-hoc tests for the PE-Valence within each stress group were performed with the Wilcoxon test. To test whether the variance in affective response and the cortisol response created by our stress manipulation is related to a bias in the updating behavior, we calculated a normalized learning rate valence bias score (valence bias score = (α_PE+_ - α_PE−_)/(α_PE+_ + α_PE−_))^21,53,54^ and correlated it with negative affect and the cortisol response using Spearman correlations. In case of the cortisol response, partial correlations controlling for time of the day were calculated to take into account circadian fluctuations of cortisol levels. To test whether the learning bias is associated with the recovery from negative affect elicited by stress (change in affective ratings post-stress T2_AFF_ – post-learning T3_AFF_), a partial correlation of the valence bias score and recovery was computed, controlling for stress-induced negative affect to take into account regression to the mean. For the whole sample, this was done using a partial Spearman correlation, while for the subsample of the social-evaluative stress group a partial Pearson correlation was computed. Data was analyzed in with the software R version 3.6.0^70^.

## References

1. Baumeister, R. F. & Leary, M. R. The need to belong: Desire for interpersonal attachments as a fundamental human motivation. Psychol. Bull. 117, 497–529 (1995).

2. Rohleder, N., Beulen, S. E., Chen, E., Wolf, J. M. & Kirschbaum, C. Stress on the dance floor: the cortisol stress response to social-evaluative threat in competitive ballroom dancers. Personal. Soc. Psychol. Bull. 33, 69–84 (2007).

3. Kirschbaum, C., Pirke, K.-M. & Hellhammer, D. H. The ‘Trier Social Stress Test’ – A tool for investigating psychobiological stress responses in a laboratory setting. Neuropsychobiology 28, 76–81 (1993).

4. Burke, P. J. Identity processes and social stress. Am. Sociol. Rev. 56, 836–849 (1991).

5. Joëls, M. & Baram, T. Z. The neuro-symphony of stress. Nat. Rev. Neurosci. 10, 459–466 (2009).

6. Gruenewald, T. L., Kemeny, M. E., Aziz, N. & Fahey, J. L. Acute threat to the social self: Shame, social self-esteem, and cortisol activity. Psychosom. Med. 66, 915–924 (2004).

7. Campbell, J. & Ehlert, U. Acute psychosocial stress: Does the emotional stress response correspond with physiological responses? Psychoneuroendocrinology 37, 1111–1134 (2012).

8. Müller-Pinzler, L. et al. Neural pathways of embarrassment and their modulation by social anxiety. Neuroimage 119, 252–261 (2015).

9. Markus, H. R. & Wurf, E. The dynamic self-concept: A social psychological perspective. Annu. Rev. Psychol. 38, 299–337 (1987).

10. Eisenberger, N. I., Inagaki, T. K., Muscatell, K. a, Byrne Haltom, K. E. & Leary, M. R. The neural sociometer: brain mechanisms underlying state self-esteem. J. Cogn. Neurosci. 23, 3448–3455 (2011).

11. Roese, N. J. & Olson, J. M. Better, stronger, faster: Self-serving judgment, affect regulation, and the optimal vigilance hypothesis. Perspect. Psychol. Sci. 2, 124–141 (2007).

12. Gotlib, I. H. & Krasnoperova, E. Biased information processing as a vulnerability factor for depression. Behav. Ther. 29, 603–617 (1998).

13. Kessler, R. C., Price, R. H. & Wortman, C. B. Social factors in psychopathology: stress, social support, and coping processes. Annu. Rev. Psychol. 36, 531–572 (1985).

14. Gloria, C. T. & Steinhardt, M. A. Relationships among positive emotions, coping, resilience and mental health. Stress Heal. 32, 145–156 (2016).

15. Bromberg-Martin, E. S. & Sharot, T. The value of beliefs. Neuron 106, 561–565 (2020).

16. Sharot, T. & Garrett, N. Forming beliefs: Why valence matters. Trends Cogn. Sci. 20, 25–33 (2016).

17. Mobius, M., Niederle, M., Niehaus, P. & Rosenblat, T. Managing self-confidence: theory and experimental evidence. (2011). doi:10.3386/w17014

18. Eil, D. & Rao, J. M. The good news-bad news effect: asymmetric processing of objective information about yourself. Am. Econ. Journal-Microeconomics 3, 114–138 (2011).

19. Korn, C. W., Prehn, K., Park, S. Q., Walter, H. & Heekeren, H. R. Positively biased processing of self-relevant social feedback. J. Neurosci. 32, 16832–16844 (2012).

20. Sharot, T., Korn, C. W. & Dolan, R. J. How unrealistic optimism is maintained in the face of reality. Nat. Neurosci. 14, 1475–1479 (2011).

21. Müller-Pinzler, L. et al. Negativity-bias in forming beliefs about own abilities. Sci. Rep. 9, 14416 (2019).

22. Ertac, S. Does self-relevance affect information processing? Experimental evidence on the response to performance and non-performance feedback. J. Econ. Behav. Organ. 80, 532–545 (2011).

23. Watabe-Uchida, M., Eshel, N. & Uchida, N. Neural circuitry of reward prediction error. Annu. Rev. Neurosci. 40, 373–394 (2017).

24. Glimcher, P. W. Understanding dopamine and reinforcement learning: The dopamine reward prediction error hypothesis. Proc. Natl. Acad. Sci. 108, 15647–15654 (2011).

25. Schultz, W., Dayan, P. & Montague, P. R. A neural substrate of prediction and reward. Science. 275, 1593–1599 (1997).

26. Adler, C. M. et al. Effects of acute metabolic stress on striatal dopamine release in healthy volunteers. Neuropsychopharmacology 22, 545–550 (2000).

27. Payer, D. et al. Corticotropin-releasing hormone and dopamine release in healthy individuals. Psychoneuroendocrinology 76, 192–196 (2017).

28. Holly, E. N. & Miczek, K. A. Ventral tegmental area dopamine revisited: effects of acute and repeated stress. Psychopharmacology. 233, 163–186 (2016).

29. Schwabe, L., Joëls, M., Roozendaal, B., Wolf, O. T. & Oitzl, M. S. Stress effects on memory: An update and integration. Neurosci. Biobehav. Rev. 36, 1740–1749 (2012).

30. van Leeuwen, J. M. C. et al. Reward-related striatal responses following stress in healthy individuals and patients with bipolar disorder. Biol. Psychiatry Cogn. Neurosci. Neuroimaging 4, 966–974 (2019).

31. Bogdan, R. & Pizzagalli, D. A. Acute stress reduces reward responsiveness: implications for depression. Biol. Psychiatry 60, 1147–54 (2006).

32. Porcelli, A. J., Lewis, A. H. & Delgado, M. R. Acute stress influences neural circuits of reward processing. Front. Neurosci. 6, 157 (2012).

33. Kumar, P. et al. Differential effects of acute stress on anticipatory and consummatory phases of reward processing. Neuroscience 266, 1–12 (2014).

34. Robinson, O. J., Overstreet, C., Charney, D. R., Vytal, K. & Grillon, C. Stress increases aversive prediction error signal in the ventral striatum. Proc. Natl. Acad. Sci. U. S. A. 110, 4129–33 (2013).

35. Garrett, N., González-Garzón, A. M., Foulkes, L., Levita, L. & Sharot, T. Updating beliefs under perceived threat. J. Neurosci. 38, 7901–7911 (2018).

36. Lighthall, N. R., Gorlick, M. A., Schoeke, A., Frank, M. J. & Mather, M. Stress modulates reinforcement learning in younger and older adults. Psychol. Aging 28, 35–46 (2013).

37. Petzold, A., Plessow, F., Goschke, T. & Kirschbaum, C. Stress reduces use of negative feedback in a feedback-based learning task. Behav. Neurosci. 124, 248–255 (2010).

38. van Leeuwen, J. M. C. et al. Increased responses of the reward circuitry to positive task feedback following acute stress in healthy controls but not in siblings of schizophrenia patients. Neuroimage 184, 547–554 (2019).

39. Nikolova, Y. S., Bogdan, R., Brigidi, B. D. & Hariri, A. R. Ventral striatum reactivity to reward and recent life stress interact to predict positive affect. Biol. Psychiatry 72, 157–163 (2012).

40. Lazarus, R. S. & Folkman, S. Stress, appraisal, and coping. (Springer Publishing Company, 1984).

41. Thoits, P. A. Stress, coping, and social support processes: where are we? What next? J. Health Soc. Behav. Spec No, 53–79 (1995).

42. Glanz, K. & Schwartz, M. D. Stress, coping, and health behavior. in Health Behavior and Health Education 211–236 (Jossey-Bass, 2008).

43. vanDellen, M. R., Campbell, W. K., Hoyle, R. H. & Bradfield, E. K. Compensating, resisting, and breaking: A meta-analytic examination of reactions to self-esteem threat. Personal. Soc. Psychol. Rev. 15, 51–74 (2011).

44. Jundt, D. K. & Hinsz, V. B. Influences of positive and negative affect on decisions involving judgmental biases. Soc. Behav. Pers. 30, 45–52 (2002).

45. Sharot, T. The optimism bias. Curr. Biol. 21, R941–R945 (2011).

46. Taylor, S. E. & Brown, J. D. Illusion and well-being: a social psychological perspective on mental health. Psychol. Bull. 103, 193–210 (1988).

47. Beck, A. T. Cognitive models of depression. in Clinical Advances in Cognitive Psychotherapy: Theory and Application (eds. Leahy, R. L. & Dowd, E. T.) 29–61 (Springer Publisher Company, 2002).

48. Korn, C. W., Sharot, T., Walter, H., Heekeren, H. R. & Dolan, R. J. Depression is related to an absence of optimistically biased belief updating about future life events. Psychol. Med. 44, 579–592 (2014).

49. Greenberg, J., Pyszczynski, T., Burling, J. & Tibbs, K. Depression, self-focused attention, and the self-serving attributional bias. Pers. Individ. Dif. 13, 959–965 (1992).

50. Hines, E. A. & Brown, G. E. A standard test for measuring the variability of blood pressure: its significance as an index of the prehypertensive state. Ann. Intern. Med. 7, 209–217 (1933).

51. Rescorla, R. A. & Wagner, A. R. A theory of Pavlovian conditioning: variations in the effectiveness of reinforcement and non reinforcement. in Classical conditioning II: current research and theory (eds. Black, A. & Prokasy, W. F.) 64–99 (Appleton-Century-Crofts, 1972).

52. Stephan, K. E., Penny, W. D., Daunizeau, J., Moran, R. J. & Friston, K. J. Bayesian model selection for group studies. Neuroimage 46, 1004–1017 (2009).

53. Niv, Y., Edlund, J. A., Dayan, P. & O’Doherty, J. P. Neural prediction errors reveal a risk-sensitive reinforcement-learning process in the human brain. J. Neurosci. 32, 551–562 (2012).

54. Palminteri, S., Lefebvre, G., Kilford, E. J. & Blakemore, S. J. Confirmation bias in human reinforcement learning: Evidence from counterfactual feedback processing. PLoS Comput. Biol. 13, e1005684 (2017).

55. Leary, M. R. Motivational and emotional aspects of the self. Annu. Rev. Psychol. 58, 317–344 (2007).

56. Hughes, B. L. & Zaki, J. The neuroscience of motivated cognition. Trends Cogn. Sci. 19, 62–64 (2015).

57. Kuzmanovic, B. & Rigoux, L. Valence-dependent belief updating: Computational validation. Front. Psychol. 8, 1087 (2017).

58. Giesler, R. B., Josephs, R. A. & Swann, W. B. Self-verification in clinical depression: The desire for negative evaluation. J. Abnorm. Psychol. 105, 358–368 (1996).

59. Pettit, J. & Joiner, T. E. Negative-feedback seeking leads to depressive symptom increases under conditions of stress. J. Psychopathol. Behav. Assess. 23, 69–74 (2001).

60. Koban, L. et al. Social anxiety is characterized by biased learning about performance and the self. Emotion 17, 1144–1155 (2017).

61. Kuzmanovic, B., Jefferson, A. & Vogeley, K. The role of the neural reward circuitry in self-referential optimistic belief updates. Neuroimage 133, 151–162 (2016).

62. Mather, M. & Lighthall, N. R. Both risk and reward are processed differently in decisions made under stress. Curr. Dir. Psychol. Sci. 21, 36–41 (2012).

63. Hines, E. A. & Brown, G. E. A standard stimulus for measuring vasomotor reactions: its application in study of hypertension. Proc. Staff Meet. Mayo Clin. 7, 332–335 (1932).

64. Brainard, D. H. The Psychophysics Toolbox. Spat. Vis. 10, 433–6 (1997).

65. Stan Development Team. RStan: the R interface to Stan, R package version 2.19.2, http://mcstan.org/. (2019).

66. Gelman, A. & Rubin, D. B. Inference from iterative simulation using multiple sequences. Stat. Sci. 7, 457–472 (1992).

67. Vehtari, A., Gelman, A. & Gabry, J. Practical Bayesian model evaluation using leave-one-out cross-validation and WAIC. Stat. Comput. 27, 1413–1432 (2017).

68. Vehtari, A., Gabry, J., Magnusson, M., Yao, Y. & Gelman, A. loo: Efficient leave-one-out cross-validation and WAIC for Bayesian models, R package version 2.1.0, https://mc-stan.org/loo. (2019).

69. Rigoux, L., Stephan, K. E., Friston, K. J. & Daunizeau, J. Bayesian model selection for group studies - revisited. Neuroimage 84, 971–85 (2014).

70. R Core Team. R: A language and environment for statistical computing. R Foundation for Statistical Computing, Vienna, Austria, http://www.r-project.org/. (2013).

