## Supplementary Information for "Self-beneficial belief updating as a coping mechanism for stress-induced negative affect"

### Supplementary Results

#### Cortisol response

*Table S1. Cortisol response - Scheirer-Ray-Hare Test*

| | *Sum of Squares* | *df* | *H* | *p* |
|---|---|---|---|---|
| Stress group | 12642 | 2 | 18.939 | < .001 |
| Time of the day | 6830 | 2 | 10.232 | .006 |
| Stress group x Time of the day | 3640 | 4 | 5.902 | .207 |
| Residuals | 35329 | 80 | | |

*Note.* Group comparison of the stress-induced cortisol response (post-stress $T2_{CORT}$ - baseline T1); *df* = degrees of freedom; *H* = test statistic; factor Stress group: social-evaluative stress (n = 29) vs. physical stress (n = 30) vs. no stress (n = 30), factor time of the day: morning vs. noon vs. afternoon.

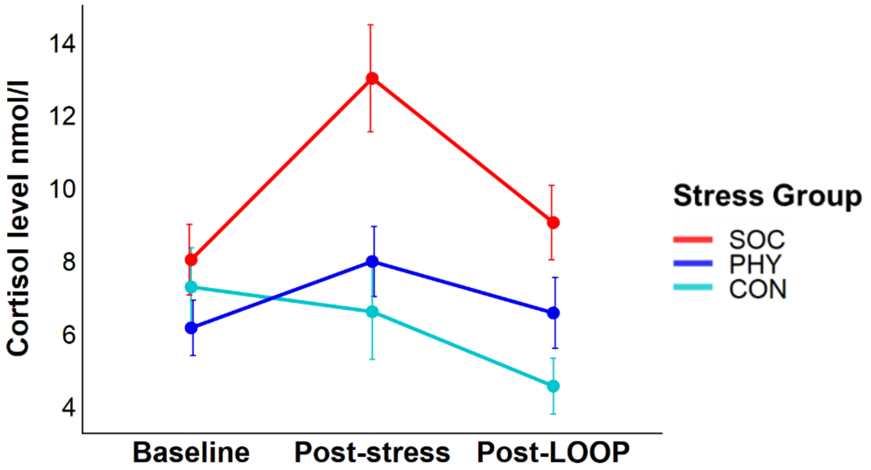


**Fig. S1**. Means and standard errors for the cortisol levels over the course of the experiment separately for the three stress groups (social-evaluative stress [n = 29] vs. physical stress [n = 30] vs. no stress [n = 30]); LOOP = Learning of own performance task.

### Negative affect ratings

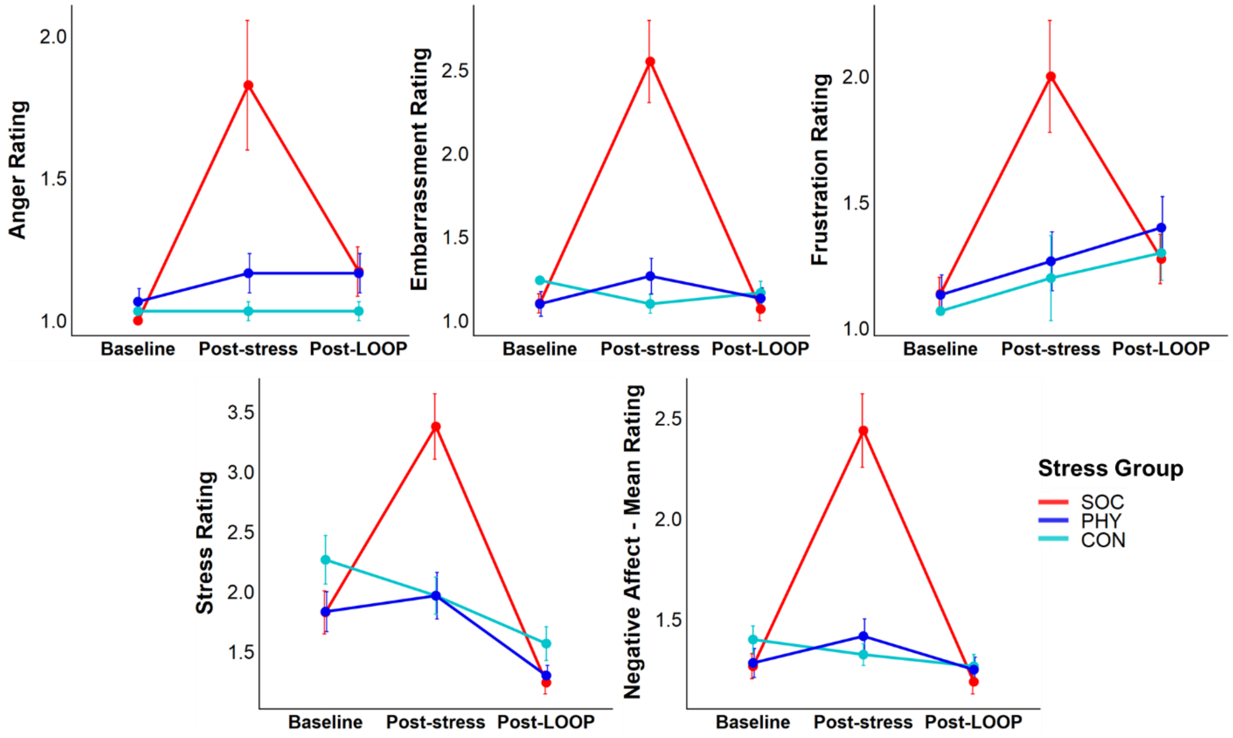


**Fig. S2**. Means and standard errors for the negative affect ratings separately for the three stress groups (SOC = social-evaluative stress [n = 29], PHY = physical stress [n = 30], CON =control [n = 30, embarrassment and frustration: n = 29 due to missing values]); LOOP = Learning of own performance task.

**Forming self-related beliefs over time - Model free behavior analysis**

*Table S2. Performance Expectation Ratings - Linear model*

| | *B [95 % CI]* | *SE* | *df* | *t* | *p* |
|---|---|---|---|---|---|
| Intercept | 42.33 [40.48; 44.19] | 0.95 | 5156 | 44.70 | < .001 |
| Ability condition | 9.59 [7.36; 11.83] | 1.13 | 86 | 8.52 | <.001 |
| Ability condition ✻ SOC vs. [PHY, CON] | -0.35 [-1.94; 1.24] | 0.80 | 86 | -0.43 | .665 |
| Ability condition ✻ PHY vs. CON | 0.79 [-1.93; 3.52] | 1.37 | 86 | 0.58 | .564 |
| Trial | -0.41 [-0.44; -0.37] | 0.02 | 5156 | -23.75 | < .001 |
| Trial ✻ SOC vs. [PHY, CON] | -0.01 [-0.04; 0.01] | 0.01 | 5156 | -0.96 | .3346 |
| Trial ✻ PHY vs. CON | 0.02 [-0.02; 0.06] | 0.02 | 5156 | 1.13 | .261 |
| Trial ✻ Ability condition | 0.80 [0.75; 0.84] | 0.02 | 5156 | 32.73 | < .001 |
| Trial ✻ Ability condition ✻ SOC vs. [PHY, CON] | 0.07 [0.04; 0.10] | 0.02 | 5156 | 4.01 | < .001 |
| Trial ✻ Ability condition ✻ PHY vs. CON | -0.06 [-0.12; -0.01] | 0.03 | 5156 | -2.15 | 0.031 |

Note. Linear mixed-effects model fit by maximum likelihood; dependent variable: performance expectation ratings; continuous variable: Trial, factor variables: Ability condition (high vs. low) and Stress group (SOC = social-evaluative stress [n = 29], PHY= physical stress [n = 30], CON = control [n = 30]) split in the contrasts SOC vs. [PHY,CON] and PHY vs. CON; *B* = unstandardized beta coefficient; *CI* = 95 % confidence interval; *SE* = standard error of B; *df* = degrees of freedom.

### Model Selection

*Table S3. Model comparison*

| *Model* | | *PSIS-LOO* | *LOO-SE* | *LOO-Diff (SE-Diff)* | *% of $\hat{k}$ > 0.7* | *No. Est. Prameters* |
|---|---|---|---|---|---|---|
| **Whole Sample** | | | | | | |
| | Unity Model | -2028.5 | 257.0 | 267.1 (52.0) | 0.09 | 3 |
| | Ability Model | -1884.4 | 247.4 | 123.0 (95.9) | 0.53 | 4 |
| | Valence Model | -1761.4 | 280.4 | | 0.17 | 4 |
| | Mean Model | -2531.9 | 219.2 | 770.5 (93.5) | 0 | 2 |
| **Social-evaluative Stress** | | | | | | |
| | Unity Model | -625.3 | 83.1 | 60.7 (21.4) | 0.17 | 3 |
| | Ability Model | -605.1 | 91.8 | 40.5 (16.4) | 0.8 | 4 |
| | Valence Model | -564.6 | 91.7 | | 0.29 | 4 |
| | Mean Model | -877.4 | 94.0 | 312.7 (40.3) | 0 | 2 |
| **Physical Stress** | | | | | | |
| | Unity Model | -840.1 | 225.7 | 107.6 (43.3) | 0 | 3 |
| | Ability Model | -782.9 | 208.8 | 50.5 (62.1) | 0.39 | 4 |
| | Valence Model | -732.5 | 247.5 | | 0.11 | 4 |
| | Mean Model | -905.5 | 181.1 | 173.1 (75.1) | 0 | 2 |
| **Control** | | | | | | |
| | Unity Model | -563.1 | 92.2 | 98.7 (19.9) | 0.11 | 3 |
| | Ability Model | -496.4 | 96.8 | 32.1 (17.3) | 0.40 | 4 |
| | Valence Model | -464.3 | 98.3 | | 0.11 | 4 |
| | Mean Model | -749.0 | 84.6 | 284.7 (35.3) | 0 | 2 |

*Note.* LOO = sum PSIS-LOO, approximate leave-one-out cross-validation (LOO) using Pareto-smoothed importance sampling (PSIS); LOO-SE = Standard error of PSIS-LOO; LOO-Diff (SE-Diff) = Difference in expected predictive accuracy (PSIS-LOO) for all models from the model with the highest PSIS-LOO (Valence Model) and standard errors of differences; percentage of $\hat{k}$ - estimated shape parameters of the generalized Pareto distribution - exceeding 0.7 (all according to Vehtari et al.[1]); No. Est. Parameters = number of estimated parameters in the model; social-evaluative stress (n = 29), physical stress (n = 30), control (n = 29).

**Posterior predictive checks: Behavioral analyses on the predicted data.**

*Table S4. Predicted Performance Expectations - Linear model*

| | *B [95 % CI]* | *SE* | *df* | *t* | *p* |
|---|---|---|---|---|---|
| Intercept | 42.56 [40.80; 44.33] | 0.90 | 5098 | 47.13 | < .001 |
| Ability condition | 9.05 [7.01; 11.08] | 1.03 | 85 | 8.82 | <.001 |
| Ability condition ✻ SOC vs. [PHY, CON] | -0.03 [-2.92; 2.86] | 1.45 | 85 | -0.02 | .983 |
| Ability condition ✻ PHY vs. CON | 1.30 [-1.56; 4.17] | 1.44 | 85 | 0.90 | .369 |
| Trial | -0.42 [-0.44; -0.41] | 0.01 | 5098 | -49.37 | < .001 |
| Trial ✻ SOC vs. [PHY, CON] | 0.00 [-0.02; 0.03] | 0.01 | 5098 | 0.40 | .690 |
| Trial ✻ PHY vs. CON | 0.03 [0.01; 0.06] | 0.01 | 5098 | 2.83 | .005 |
| Trial ✻ Ability condition | 0.84 [0.82; 0.87] | 0.01 | 5098 | 69.66 | < .001 |
| Trial ✻ Ability condition ✻ SOC vs. [PHY, CON] | 0.07 [0.03; 0.10] | 0.02 | 5098 | 3.93 | < .001 |
| Trial ✻ Ability condition ✻ PHY vs. CON | -0.10 [-0.13; -0.07] | 0.02 | 5098 | -5.85 | < .001 |

Note. Linear mixed-effects model fit by maximum likelihood; dependent variable: performance expectations predicted by winning model; continuous variable: Trial, factor variables: Ability condition (high vs. low) and Stress group (SOC = social-evaluative stress [n = 29], PHY= physical stress [n = 30], CON = control [n = 29]) split in the contrasts SOC vs. [PHY,CON] and PHY vs. CON; *B* = unstandardized beta coefficient; *CI* = 95 % confidence interval; *SE* = standard error of B; *df* = degrees of freedom.

**Learning parameters.**

**Group comparison of learning rates**

*Table S5. Learning rates - Linear model*

| | *B [95 % CI]* | *SE* | *b* | *df* | *t* | *p* |
|---|---|---|---|---|---|---|
| Intercept | 0.091 [0.079; 0.105] | 0.007 | | 85 | 13.412 | < 0.001 |
| PE-Valence | -0.013 [-0.020; -0.006] | 0.004 | -0.178 | 85 | -3.596 | .0005 |
| SOC vs. [PHY, CON] | 0.015 [-0.004; 0.034] | 0.010 | 0.138 | 85 | 1.500 | .1373 |
| PHY vs. CON | -0.007 [-0.023; 0.010] | 0.008 | -0.074 | 85 | -0.798 | .4272 |
| PE-Valence ✻ SOC vs. [PHY, CON] | 0.012 [0.002; 0.022] | 0.005 | 0.114 | 85 | 2.303 | .0237 |
| PE-Valence ✻ PHY vs. CON | -0.003 [-0.012; 0.006] | 0.004 | -0.036 | 85 | -0.724 | .4711 |

*Note.* Linear mixed-effects model fit by maximum likelihood; dependent variable: learning rates derived from the valence model; learning rates for positive and negative prediction errors (PE, within subject factor PE-Valence); Stress group (SOC = social-evaluative stress [n = 29], PHY= physical stress [n = 30], CON = control [n = 29]) split in the contrasts SOC vs. [PHY,CON] and PHY vs. CON; *B* = unstandardized beta coefficient; *CI* = 95 % confidence interval; *SE* = standard error of *B*; *b* = standardized beta coefficient; *df* = degrees of freedom.

**Associations of valence bias score with stress response.**

As was to be expected, both measured components of the stress response, i.e. change in negative affect (ΔAFF, post-stress $T2_{AFF}$ - baseline $T1_{AFF}$) and the cortisol response (ΔCORT, post-stress $T2_{CORT}$ - baseline $T1_{CORT}$) share common variance ($\rho_{\Delta AFF,\Delta CORT}$ = .31, *p* = .003). In order to test the effect of one component on the valence bias score (BIAS, $(\alpha_{PE+} - \alpha_{PE-})/(\alpha_{PE+} + \alpha_{PE-})$) independently of the other, partial correlations were calculated additionally. The change in negative affect could only trend wise predict a learning bias independent of the cortisol response ($\rho_{BIAS,\Delta AFF|\Delta CORT}$ = .18, *p* = .097), the effect of the cortisol response on the learning bias remained significant when controlling for the negative affect response and time of the day (TIME, $\rho_{BIAS,\Delta CORT|\Delta AFF,TIME}$ = .23, *p* = .035). Within the subsamples of the three stress groups all correlations between change in negative affect/ cortisol response and the valence bias score are not significant (Table S6).

*Table S6. Spearman correlations for change in negative affect and cortisol stress reaction with the valence bias score*

| | | **Social Stress** | | **Physical Stress** | | **Control** | |
|---|---|---|---|---|---|---|---|
| | | **ΔAFF** | **ΔCORT\| TIME** | **ΔAFF** | **ΔCORT\| TIME** | **ΔAFF** | **ΔCORT\| TIME** |
| **Valence Bias Score** | *ρ* | .03 | .05 | .34 | .32 | -.09 | .18 |
| | *p* | .865 | .807 | .063 | .092 | .660 | .360 |
| | *n* | 29 | 29 | 30 | 30 | 29 | 29 |

*Note*. Valence bias score = $(\alpha_{PE+} - \alpha_{PE-})/(\alpha_{PE+} + \alpha_{PE-})$, ΔAFF = change in negative affect (post-stress $T2_{AFF}$ - baseline $T1_{AFF}$), ΔCORT = Cortisol change (post-stress $T2_{CORT}$ - baseline $T1_{CORT}$) controlled for TIME = time of the day (morning vs. noon vs. afternoon); *ρ* = Spearman's Rho.

*Table S7. Partial correlations of recovery from negative affect with the valence bias score controlled for initial change in negative affect*

| | **Recovery from negative affect\| ΔAFF** | | | | | | | | |
|---|---|---|---|---|---|---|---|---|---|
| | **Social Stress** | | | **Physical Stress** | | | **Control** | | |
| | *r* | *p* | *n* | *ρ* | *p* | *n* | *ρ* | *p* | *n* |
| **Valence Bias Score** | .382 | .045 | 29 | .232 | .226 | 30 | .006 | .977 | 29 |

*Note*. Recovery from negative affect = post-stress $T2_{AFF}$ - post-learning $T3_{AFF}$,
Valence bias score = $(\alpha_{PE+} - \alpha_{PE-})/(\alpha_{PE+} + \alpha_{PE-})$, ΔAFF = change in negative affect (post-stress $T2_{AFF}$ - baseline $T1_{AFF}$); *r* = Pearson's *r*; *ρ* = Spearman's Rho.

*Table S8. Partial correlations of cortisol recovery with the valence bias score controlled for initial change in cortisol*

| | Cortisol recovery \| ΔCORT | | | | | | | | | | | |
|---|---|---|---|---|---|---|---|---|---|---|---|---|
| | **Social Stress** | | | **Physical Stress** | | | **Control** | | | **Whole sample** | | |
| | *r* | *p* | *n* | *ρ* | *p* | *n* | *ρ* | *p* | *n* | *ρ* | *p* | *n* |
| **Valence Bias Score** | -.159 | .420 | 29 | -.008 | .967 | 30 | -.094 | .635 | 29 | -.012 | .913 | 88 |

*Note*. Cortisol recovery = post-stress $T2_{CORT}$ - post-learning $T3_{CORT}$; Valence bias score = $(\alpha_{PE+} - \alpha_{PE-})/(\alpha_{PE+} + \alpha_{PE-})$; ΔCORT = stress-induced cortisol change (post-stress $T2_{CORT}$ - baseline $T1_{CORT}$); *r* = Pearson's *r*; *ρ* = Spearman’s Rho.

### Supplementary Tables

*Table S9 a. Sample characteristics*

| | Social Stress | | | Physical Stress | | | Control | | | Test | |
|---|---|---|---|---|---|---|---|---|---|---|---|
| | *M* | *Md* | *SD* | *M* | *Md* | *SD* | *M* | *Md* | *SD* | *H(2)* | *p* |
| **Age** | 22.9 | 23 | 2.76 | 22.5 | 23 | 1.94 | 22.3 | 22 | 3.00 | 1.47 | .480 |
| **Self-esteem** | 6.44 | 6.75 | 1.02 | 6.3 | 6.42 | 0.94 | 6.02 | 6.25 | 0.93 | 5.03 | .080 |
| **SIAS** | 1.91 | 1.9 | 0.51 | 1.96 | 1.92 | 0.31 | 2.02 | 2 | 0.6 | 1.23 | .540 |
| **Cortisol baseline** | 8.04 | 7.07 | 5.22 | 6.17 | 4.88 | 4.17 | 7.3 | 5.09 | 5.89 | 1.74 | .419 |
| **Affective state baseline** | 1.27 | 1.25 | 0.33 | 1.28 | 1.25 | 0.39 | 1.4 | 1.25 | 0.39 | 3.21 | .201 |

*Note.* Sample characteristics for the three stress groups. *M* = mean; *Md* = median; *SD* = standard deviation; self-esteem assessed via averaged scores of the Self-Description Questionnaire (SDQ-III); SIAS = averaged score on the Social Interaction Anxiety Scale; *H* = Kruskal-Wallis Chi-squared.

*Table S9 b. Sample characteristics*

| | | Social Stress | Physical Stress | Control | Test | |
|---|---|---|---|---|---|---|
| | | | | | *H* | *p* |
| **Gender** | **female** | 21 | 20 | 20 | | |
| | **male** | 8 | 10 | 10 | 0.3 (*df*=2) | .861 |
| **Time of day** | **morning** | 10 | 10 | 10 | | |
| | **noon** | 11 | 10 | 8 | | |
| | **afternoon** | 8 | 10 | 12 | 1.27 (*df*=4) | .867 |

*Note*. Frequency distribution for gender and time of day of the measurement for the three stress groups; *H* = Pearson's Chi-squared test statistic

### References

1. Vehtari, A., Gelman, A. & Gabry, J. Practical Bayesian model evaluation using leave-one-out cross-validation and WAIC. *Stat. Comput.* **27**, 1413–1432 (2017).
